# Arrhythmic gut microbiome signatures for risk profiling of Type-2 Diabetes

**DOI:** 10.1101/2019.12.27.889865

**Authors:** Sandra Reitmeier, Silke Kießling, Thomas Clavel, Markus List, Eduardo L. Almeida, Tarini S. Ghosh, Klaus Neuhaus, Harald Grallert, Martina Troll, Wolfgang Rathmann, Birgit Linkohr, Andre Franke, Caroline I. Le Roy, Jordana T. Bell, Tim Spector, Jan Baumbach, Peter W. O’Toole, Annette Peters, Dirk Haller

## Abstract

To combat the epidemic increase in Type-2-Diabetes (T2D), risk factors need to be identified. Diet, lifestyle and the gut microbiome are among the most important factors affecting metabolic health. We demonstrate in 1,976 subjects of a prospective population cohort that specific gut microbiota members show diurnal oscillations in their relative abundance and we identified 13 taxa with disrupted rhythmicity in T2D. Prediction models based on this signature classified T2D with an area under the curve of 73%. BMI as microbiota-independent risk marker further improved diagnostic classification of T2D. The validity of this arrhythmic risk signature to predict T2D was confirmed in 699 KORA subjects five years after initial sampling. Shotgun metagenomic analysis linked 26 pathways associated with xenobiotic, amino acid, fatty acid, and taurine metabolism to the diurnal oscillation of gut bacteria. In summary, we determined a cohort-specific risk pattern of arrhythmic taxa which significantly contributes to the classification and prediction of T2D, highlighting the importance of circadian rhythmicity of the microbiome in targeting metabolic human diseases.

---

Increasing evidence links the human gut microbiome to metabolic health (1), and altered microbial profiles are associated with obesity, insulin resistance, and Type-2-Diabetes (T2D) (2–9). Population-based studies highlighted a significant degree of variability in inter-individual microbiome differences (10, 11), regional effects (12), and drug-associated changes in the gut microbiome (13, 14), which complicates the identification of disease-related microbial risk factors. Despite extensive investigations of the role of the gut microbiome in metabolic diseases, especially T2D, there is still no consensus on disease-related bacterial taxa with diagnostic relevance.

The circadian clock, which synchronizes daily food intake behaviour and metabolism with the day and night cycle (15), has recently been proposed to influence microbial homeostasis (16). Daytime-dependent fluctuations were identified in both the oral and faecal microbiota (16, 17). Circadian rhythms in gut microbiota composition and function are sensitive to diet and feeding patterns in murine models (16, 18). Although diet-induced obesity dampens cyclic microbial fluctuations in rodents (18), and epidemiological studies continue to show associations between circadian clock dysfunction due to modern lifestyle and T2D (reviewed in (19)), the clinical relevance of daily microbial oscillations in human diseases is unclear and requires confirmation in human population studies.

To understand whether circadian rhythms affect microbiome features related to the onset and progression of metabolic diseases, we considered the daytime of stool sampling in microbiota analysis of a regionally confined prospective cohort (KORA). We provide the first demonstration of robust diurnal oscillations in faecal microbiota composition analysed in a large-scale population and propose a functional role of circadian rhythmicity of the gut microbiome in regulating metabolic health.

## Results

### Microbiota profiling of the population-based cohort KORA

KORA is a prospective cohort in the region of Augsburg (Germany) designed to understand the role of genetic, life-style and environmental factors in disease progression including metabolic disease (**Supplementary Table 1**). As part of the third follow-up of the S4 KORA cohort, stool was sampled from 1,976 individuals in 2013, for whom we performed high-throughput 16S rRNA gene amplicon sequencing (**Supplementary Table 2-3; Extended Data Table 1**). Comparing individual microbiota compositions confirmed diverse ecosystems dominated by the two major phyla *Firmicutes* and *Bacteroidetes* (cumulative mean relative abundance, 91%) (**Fig. 1 A, B**). In comparison to other studies, compositional variations were marginally affected by geography (0.9%), since KORA is restricted to a single city and its close surrounding (12, 20, 21) (**Figure 1 C).** The cohort was characterized by an average richness of 348 ±77 operational taxonomic units (OTUs) and 118± 37 Shannon effective counts **(Fig. 1 D**).

**Figure 1.**
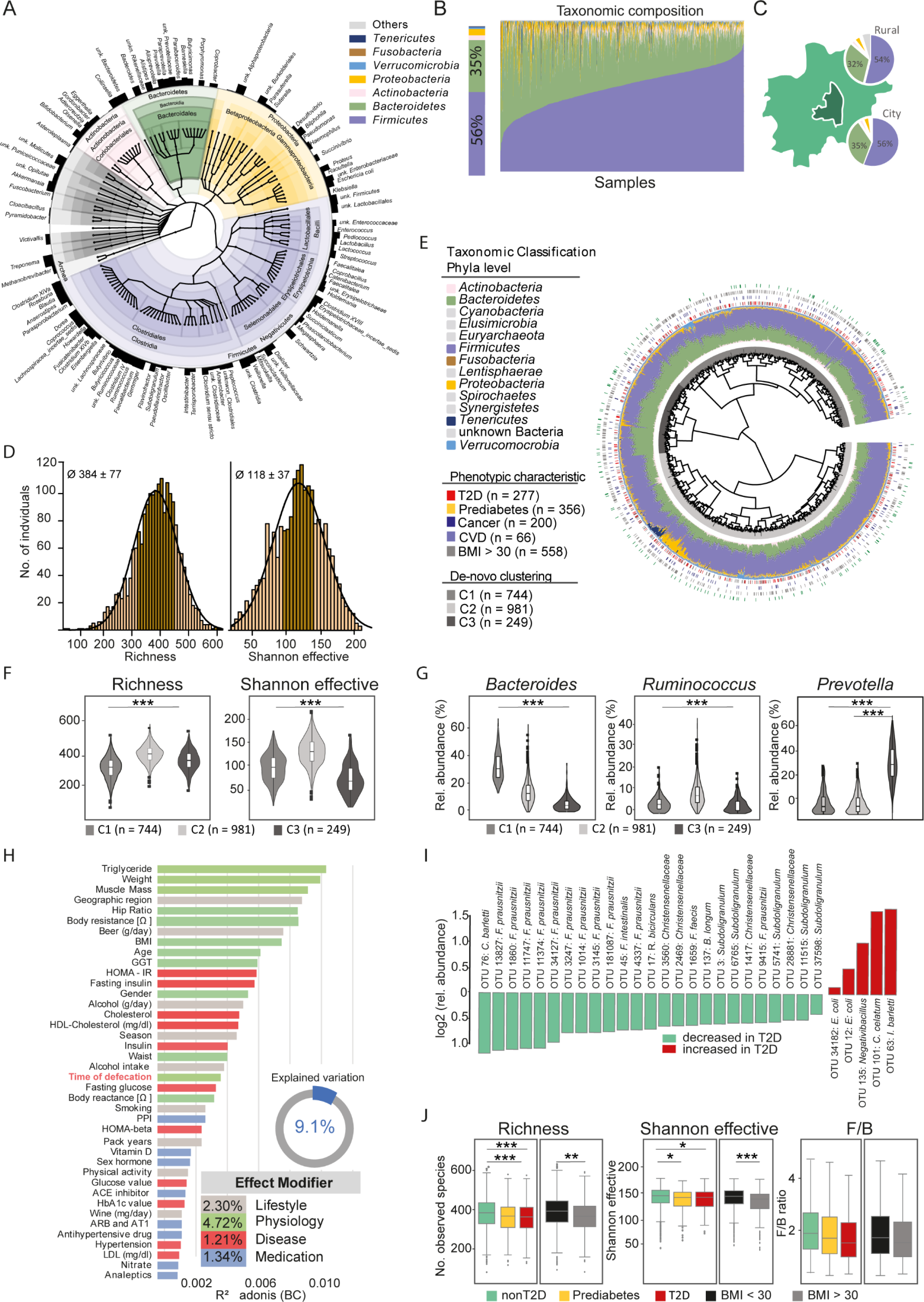
Microbiota profiling of a cross-sectional population-based cohort. **1A:** Taxonomic tree of the gut microbiota in 1,976 KORA subjects. Colors indicate phyla. Taxonomic ranks are from kingdom (center) to genera indicated by the individual branches. Black bars indicate the prevalence of each genus, the name of which are shown if found in > 10% of individuals (Supplementary Table 1). **1B:** Relative abundances of phyla across the whole cohort. Samples are ordered according to increasing relative abundances of *Firmicutes*. Colors are as in Fig.1A. **1C:** Geographical map of the city of Augsburg and its rural area. Subjects are grouped according to their place of residence. The pie charts show taxonomic distributions at the phylum level in individuals living outside (rural) or in the city. **1D:** *Alpha*-diversity of the faecal microbiota in KORA. Richness (left; 384 ±77) and Shannon effective counts (right; 118 ±37), which were both not normally distributed across the whole cohort (Shapiro test < 0.05). **1E:** *Beta*-diversity of the faecal microbiota in KORA. The dendrogram shows similarities between microbiota profiles based on generalized UniFrac distances between 1,976 subjects represented by individual branches. Unsupervised hierarchical clustering identified three main clusters of individuals (grey-scale next to branches). Individual taxonomic composition at the phylum level is shown as stacked bar plots around the dendrogram and follows the color code as in panel A. Bars in the outer part of the figure indicate disease status: first ring, diabetes status (red, T2D; grey, Prediabetes; no color, nonT2D); second ring, cancer (blue); third ring, obesity (grey, BMI ≥ 30); fourth ring, cardiovascular diseases (green). **1F:** Differences in *alpha*-diversity between the *de-novo* clusters from panel E. **1G:** Differences in relative abundances of the three genera *Bacteroides*, *Ruminococcus* and *Prevotella* for the three microbiota clusters as in panel E. *** P < 1·10^−5^. **1H**: Explained variations in faecal microbiota composition by covariates. All variables shown had a significant influence (P ≤ 0.05) displayed as proportions of explained variations based on R^2^ calculated by multivariate analysis of Bray-Curtis dissimilarity. **1I:** OTUs (N = 30) with significantly different relative abundances between T2D and nonT2D subjects (Supplementary Table 2). **1J:** Significant differences in *alpha*-diversity and *Firmicutes* to *Bacteroidetes* ratios depending on metabolic health (red, T2D; yellow, prediabetes; green, nonT2D; grey, BMI ≥ 30; black, BMI < 30).

Unsupervised analysis based on generalized UniFrac distances identified three faecal microbiota clusters (C1, N = 744; C2, N = 981; C3, N = 249) similar to previously reported enterotypes (20) (**Fig. 1 E-G**). Individuals with obesity (BMI ≥ 30, N = 558), T2D (N = 277) and pre-diabetic conditions classified according to their oral glucose tolerance (N = 356, WHO criteria), cancer (N = 200), and cardiovascular disease (CVD, N = 66) were evenly distributed across these clusters (**Fig. 1 E**). Individuals in C1 had the lowest microbiota richness and showed significantly higher relative abundances of *Bacteroides*. The most diverse cluster C2 (highest number of subjects) was dominated by members of the genus *Ruminococcus*, whilst *Prevotella* dominated in C3. Multivariate analysis of metadata co-varying with the faecal microbiota profiles identified 40 of 113 features related to physiology (*e.g.* blood triglyceride levels, body weight, muscle mass, time of defecation), lifestyle and environment (geographical region, beer/alcohol consumption, seasons), disease-associated parameters (mostly related to glucose metabolism), and medication, collectively explaining 9.1% of variability (**Fig. 1 H**). In this study, we focused on individuals with risk factors for metabolic disorders, including obese, prediabetic and T2D subjects. Significantly different relative abundances were identified for 30 OTUs in T2D subjects (N = 277) *vs.* all others (N = 1,270) (**Fig. 1 I**; **Extended Data Table 2**). Species richness and *alpha*-diversity were lower in individuals with T2D and obesity (BMI ≥30), while *Firmicutes* to *Bacteroidetes* ratios remained unchanged compared to healthy subjects (**Fig.1 J).**

### Diurnal rhythms in faecal microbiota composition

Time of defecation was among the most significant factors explaining inter-individual variabilities in microbiota structure (**Fig. 1 H**). Thus, diurnal rhythmicity of faecal microbiota profiles was studied in all 1,943 subjects for whom time at sampling was available (**Fig. 2**). Community diversity (both species richness and Shannon effective counts) fluctuated significantly throughout the day (**Fig. 2 A**). Diurnal rhythmicity was also evident in relative abundances of the two most dominant phyla, which oscillated in antiphase. *Bacteroidetes* showed 6% higher mean relative abundance at night, while the phylum *Firmicutes* was higher during the day. Since more than 70% of all samples were collected at morning hours between 5 and 11 am, we re-analysed the data using matched sample sizes, and thereby confirmed initial results including all subjects (**Supplementary Fig. 1 A**).

**Figure 2.**
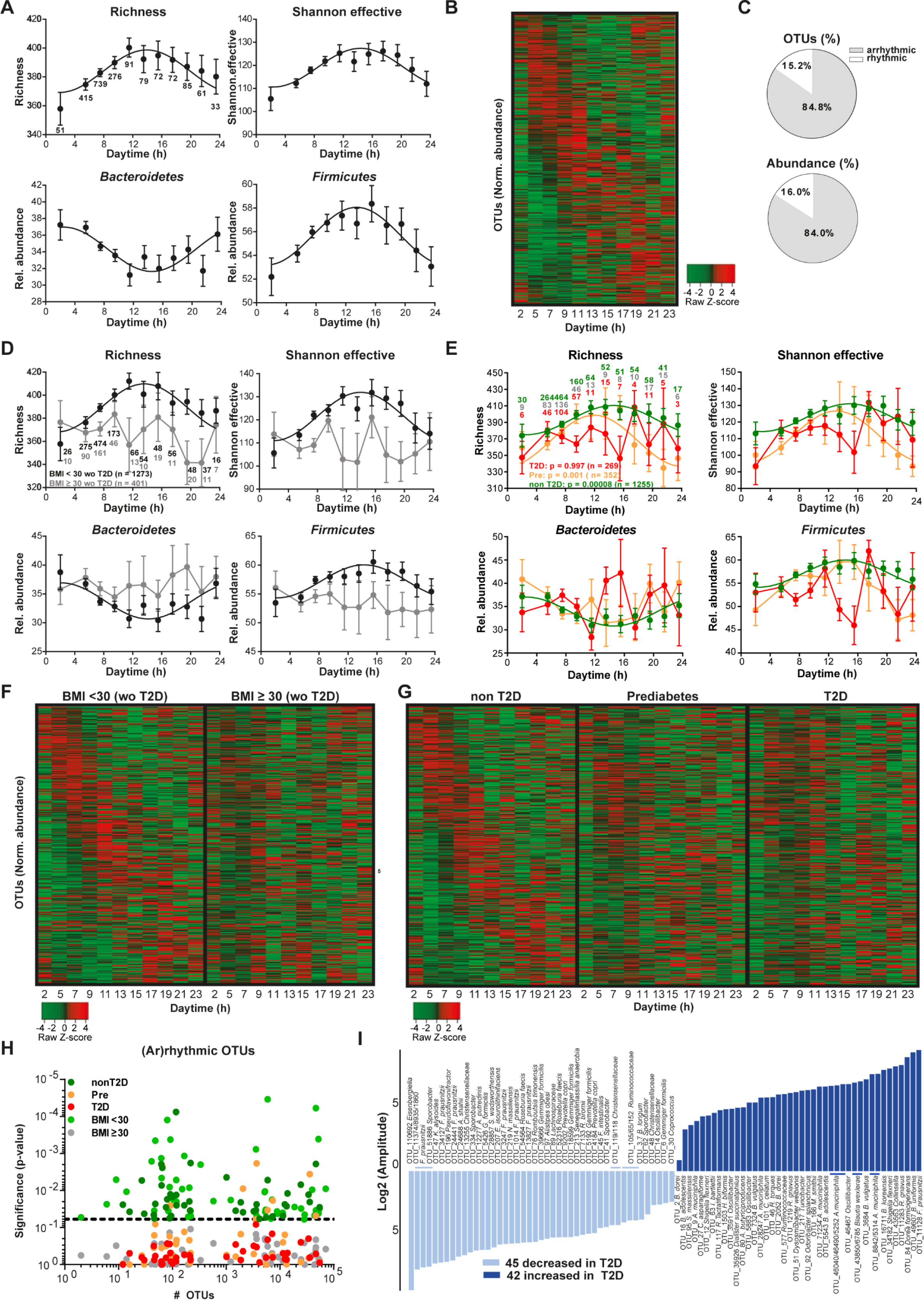
Diurnal rhythms in the human gut microbiota. **2A:** Diurnal profiles of *alpha*-diversity and of relative abundances of the phyla *Bacteroidetes* and *Firmicutes* in 1,943 subjects. Significant rhythms are illustrated with fitted cosine-wave curves (cosine-wave regression, P ≤ 0.05). **2B:** Heatmap depicting the overall phase relationship and periodicity of OTUs (N = 422; mean relative abundance >0.1%; prevalence >10% individuals) ordered by their cosine-wave peak phase according to time of day and normalized to the peak of each OTU. **2C:** Amount of rhythmic (white) and arrhythmic (grey) OTUs and their relative abundance in percent (compare to panel B). **2D:** Diurnal profiles of *alpha*-diversity and of relative abundances of the phyla *Bacteroidetes* and *Firmicutes* of subjects without diabetes (wo T2D), either shown for a BMI < 30 (black, N = 1,273) or a BMI ≥ 30 (grey, N = 401). **2E:** As panel D, but of subjects with diabetes (T2D, red; N = 269), with Prediabetes (Pre, orange; N = 352, orange) or without diabetes (non T2D, green; N = 1,254). Significant rhythms (cosine-wave regression, P ≤ 0.05) are illustrated with fitted cosine-wave curves; data points connected by straight lines indicate no significant cosine fit curves (P > 0.05) and thus no rhythmicity. **2F:** Heatmap of the normalized daytime-dependent relative abundance of OTUs based on 422 OTUs (see panel A). Data are normalized to the peak of each OTU and ordered by the peak phase according to subjects without diabetes but a BMI < 30 (left) or ≥ 30 right). **2G:** As panel F, but of subjects without diabetes (left, nonT2D) with prediabetes (middle) or with diabetes (right, T2D). **2H:** OTUs show diurnal rhythms in control groups (dark green, nonT2D, N = 1,254; light green, BMI < 30, N = 546), but are arrhythmic in disease stages like prediabetes (orange, Pre, N = 352), diabetes (red, T2D, N = 269), or obesity (grey, BMI ≥ 30, N = 1,396). Significance of rhythmicity (y-axis) is indicated by P-values below the dashed line (P ≤ 0.05; cosine-wave regression). **2I:** Amplitude of OTUs rhythmic in healthy controls but arrhythmic in T2D. Of 422 OTUs (compare to panel G), 87 OTUs oscillated in controls only. These OTUs are ordered according to their fluctuation amplitude in healthy controls: light blue, decreased; deep blue, increased relative abundance in T2D (Supplementary Table 2).

After removal of OTUs low in mean relative abundance (<0.1%) and prevalence (<10% subjects), a heatmap of the remaining 422 OTUs illustrated the heterogeneous distribution of their peak relative abundances, ranging from early day to late night, suggesting that the microbiota at different times of the day are dominated by different microbial taxa (**Fig. 2 B; Extended Table 3**). 15.2% of the OTUs were rhythmic (rOTUs) according to cosine-wave regression analysis (**Fig. 2 C**). Similar proportions of rOTUs were identified using other non-parametric (12.3%) and parametric methods (13.5%; **Supplementary Fig. 1 B**), demonstrating validity of the analysis.

### Microbial oscillations are disrupted in obesity and T2D

Robust daily oscillations in *alpha*-diversity, phyla and molecular species were observed in subjects with BMI < 30 (N = 1,273 without T2D) and subjects without T2D (nonT2D; N = 1,255) (**Fig. 2 D-G; Supplementary Fig. 2 A-D; Supplementary Fig. 3 A-D**). In contrast, rhythmicity in *alpha*-diversity and phylum proportions (*Bacteroidetes* and *Firmicutes*) were completely absent in subjects with either a BMI ≥ 30 excluding all T2D cases (N = 401) (**Fig. 2 D)** or T2D regardless of BMI (N = 269) (**Fig. 2 E).** Heatmaps showing peak relative abundances of OTUs confirmed the disruption of rhythmicity in subjects with BMI ≥ 30 (**Fig. 2 F; Supplementary Fig. 3 D)** and T2D (**Fig. 2 G, Supplementary Fig. 2 B).** All 10.4% OTUs that oscillated in nonT2D subjects lost rhythmicity in subjects with T2D (**Fig. 2 G, Supplementary Fig. 2 A).** To account for the difference in sample size between subject groups (T2D, N = 269; nonT2D, N = 1255), the circadian analysis was tested using 10 different randomly selected and sample size-matched groups (**Supplementary Fig. 2 D**). Loss of diurnal oscillations in diabetic and obese subjects was well reflected in the relative abundances of single OTUs (**Fig. 2 F, G**). OTUs with disrupted rhythmicity in T2D were to a large extent (>60%) not shared with arrhythmic OTUs in obese individuals, indicating a BMI-independent loss of rhythmicity in T2D (compare **Fig. 2F, G; Supplementary Fig. 2 B, C**). Interestingly, intermediate phenotypes were noted in prediabetic subjects (N = 352) with a loss of rhythmicity for the two phyla but not alpha-diversity (**Fig. 2 E**). In prediabetes, the proportion of rOTUs was reduced from 10.4% to 7.6% (**Fig. 2 G; Extended Data Fig. 2 A**). Similar results were obtained using JTK_CYCLE or harmonic cosine-wave regression, demonstrating robustness of the findings (**Supplementary Fig. 2 A**). We identified 87 OTUs that oscillated in controls but lacked rhythmicity in T2D. They belonged to the genera *Akkermansia, Bacteroides*, *Bifidobacterium*, *Blautia*, *Clostridium, Coprococcus, Dorea, Prevotella, Roseburia* and *Ruminococcus* (**Fig. 2 H, I; Extended Data Table 2**), which is in line with recently published data describing oscillations in two subjects (16). Altogether, these population-based findings clearly indicate that rhythmicity of the faecal microbiota is disrupted in subjects with obesity and T2D.

### Classification and prediction of T2D using arrhythmic microbial signatures

We then sought to identify diagnostic biomarkers for T2D development using microbiota profiles of 1,340 subjects sampled in 2013 as training data and another 699 subjects for whom matched samples at the 5-year follow-up (2018) were available as independent test data (**Fig. 3 A, Supplementary Table 3**). Among the 87 arrhythmic (**Fig. 2 H, I**) we selected 13 OTUs with differential 24-h time-of-day patterns using the Detection of Differential Rhythmicity (DODR) R packages (22) (s-arOTUs) (**Fig. 3 B; Extended Data Table 2**), which overlap with the 30 differentially abundant OTUs detected in the whole cohort (**Fig. 1 I).** We trained a generalized linear model with these 13 s-arOTUs to classify T2D. The model performed significantly better than an equal number of randomly selected control OTUs (rndOTUs) not rhythmic in either of the groups (repeated 100-times, mean area under the curve AUC = 0.79 vs. 0.59 P-value for 100 permutations = 2.03·10^−8^) (**Fig. 3 C; Supplementary Fig. 2 E**).

**Figure 3.**
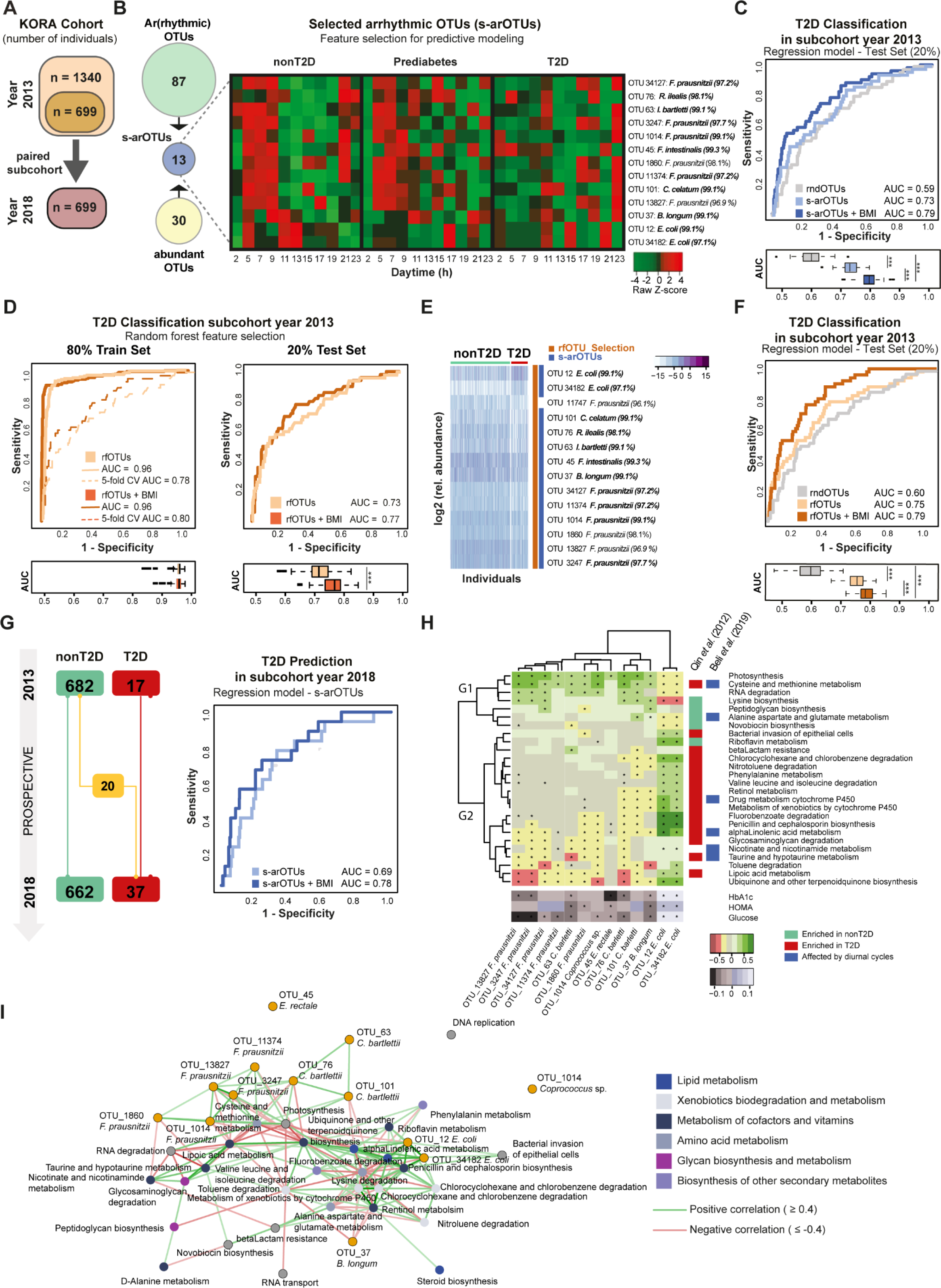
Arrhythmic microbial signature for classification and prediction of T2D and T2D associated functional pathways. **3A:** Number of KORA subjects of the prospective sub-cohort with samples from both year 2013 and 2018 (N = 699; **Extended Data Table 3**). **3B: Left panel**, Among the 87 arOTUs (green circle), which oscillated in controls (see Fig. 2F, G) but are arrhythmic in subjects with T2D or BMI ≥ 30, 13 OTUs (blue circle) showed differential rhythmicity based on DODR analysis and overlapped with the previously defined 30 OTUs (yellow circle) with a significantly different relative abundance (see Fig. 1I). These 13 selected arrhythmic OTUs (s-arOTUs) were used for further generalized linear model (Supplementary Table 2). **Right panel**, heatmap representing the relative abundance of the s-arOTUs according to the time of day. **3C:** Curves of receiver operating characteristics (ROC) for classification of T2D in an independent test set. The generalized linear model is based on 13 s-arOTUs +/− BMI (blue curves) as well as on 100 randomly selected sets of 13 OTUs (grey curve). The distribution of AUCs are shown by boxplots and are significantly different between the types of models. **3D:** ROC curve for a random forest model using a training set (train set) of 80% of the data (dashed lines in the left panel) as well as using a test set with the remaining 20% of the data (ROC curves in the right panel). The mean AUC over 100 random data splits is shown. The boxplots below the curve panels show the distribution of AUCs across all generated models for the corresponding training and test sets, respectively. **3E:** Heatmap showing the log-transformed relative abundance of the OTUs (y-axis) selected by the random forest model. Individuals (x-axis) are grouped according to their T2D status (Supplementary Table 3). Taxonomic classification of OTUs is shown by their species names and 16S rRNA gene sequence identities (%); bold letters indicate ≥97% identity (proxy for species level). To the right, overlap representation of the 14 rfOTUs (orange, according to Fig. 3C) and the 13 s-arOTUs (dark blue). **3F:** ROC curve for classification of T2D in an independent test set consisting of 20% of the data. The generalized linear model is based on 14 selected OTUs +/− BMI (rfOTUs, orange curves) as well as on the average performance of 100 random sets of 14 OTUs (rndOTUs, grey curve). The distribution of AUCs of all models is shown by boxplots and differed significantly between the three model types. **3G**: Predictive analysis of T2D in the paired KORA sub-cohort (samples from 2013 and 2018). The prediction of incident T2D cases (iT2D) is based on the 13 s-arOTUs +/− BMI. Baseline data is from nonT2D individuals from 2013. Endpoint is iT2D in 2018 (N = 20). **3H**: The heatmap shows the Spearman correlations for 26 disease-predictive microbial pathways within the 13 OTUs becoming arrhythmic in T2D (across the 100 shotgun sequenced samples). Only pathways that were significantly correlated with at least one of the 13 arrhythmic OTUs are shown. The heatmap on the bottom left shows the association pattern for the 13 OTUs with clinical markers characterizing T2D (across the entire cohort). Significant associations of both heatmaps were corrected using the Benjamini-Hochberg procedure. Corrected P ≤ 0.1 are indicated with * in each field. On the right, a one-dimensional heat-plot shows relative representations of each of the previous pathways observed in the cohort of Qin et al. (2012) (4), either enriched in T2D (red) or enriched in nonT2D (green). Furthermore, pathways predicted to be influenced by diurnal cycles in Beli et al. (2019) (24) are indicated in blue if affected. **3I**: Correlation network of HbA1c associated pathways in relation to selected T2D markers (s-arOTUs; orange circles). Green and red lines indicate a positive and negative association, respectively. The thickness of lines is proportional to correlation coefficients. The color of circles refers to functional classes as indicated.

As a complementary and hypothesis-free approach for identifying T2D biomarkers, we also trained a random forest (RF) model, in which a 5-fold cross validation was applied to 80% of the data, while the remaining 20% were used to assess performance. We repeated this random split 100 times and identified 63 out of 425 OTUs that were consistently selected as predictive with a mean AUC of 0.73 on the test set (**Fig. 3D**). BMI as an additional variable in the model, reduced the number of selected OTUs to 14 (rfOTUs) with significant differences in relative abundance (mean AUC = 0.77, **Fig. 3 D, E, Extended Data Table 2**). This signature included *Bifidobacterium longum* (OTU 37)*, Clostridium celatum* (OTU 101*), Intestinibacter bartletti* (OTU 63)*, Romboutsia ilealis* (OTU 76), and several taxa closely related to *Faecalibacterium prausnitzii* (OTU 34127, OTU 3247, OTU 1860, OTU 11374, OTU 1014) and *Escherichia coli* (OTU 12, OTU 34182). A model trained on the outcome obesity was not able to differentiate T2D (mean AUC = 0.68), suggesting that the selected 14 rfOTUs are not merely surrogates of the confounding variable BMI. In reverse, the selected 14rfOTUs failed to differentiate obesity (mean AUC = 0.58; **Supplementary Fig. 3 E, F**). Strikingly, 13 of these 14 rfOTUs are shared with s-arOTUs (**Fig. 3 B, E**). A generalized linear model using the selected rfOTUs and BMI (rfOTUs+BMI) classified T2D with an AUC of 0.79, performing significantly better than a set of 14 randomly picked OTUs (rndOTUs, AUC = 0.60; repeated 100-times, **Fig. 3 F**). Due to a comparable performance of both models and the extensive overlap of OTUs, all 13 s-arOTUs were used for further analysis.

In agreement with previous results (13, 14), metformin intake (MET) significantly affected T2D classification (+MET T2D AUC = 0.87 vs. -MET T2D AUC = 0.60; **Supplementary Fig. 4 A**), but a risk signature based on +/-MET intake was not able to classify T2D in the KORA cohort (AUC = 0.60; **Supplementary Fig. 4 B**). Although MET was found to synchronize peripheral circadian clocks (23), indicating that MET may directly interfere with the circadian analysis, MET did not change rhythmicity nor the overall percentage of rOTUs in subjects with T2D in our study (**Supplementary Fig. 4 C, D).** Moreover, none of the rOTUs identified in +/− MET-T2D overlap with the 13 selected arrhythmic OTUs (s-arOTUs) used for the classification of T2D (**Supplementary Fig. 4 E**). Consequently, MET did not affect the predictability of T2D based on the s-arOTU signature.

We then applied the 13 s-arOTU to predict T2D in the prospective arm of KORA, which included 699 paired samples from 2013 and 2018 with 17 persisting T2D (pT2D) and 20 newly incident T2D (iT2D) cases (**Fig. 3 G, Supplementary Table 3**). Both sub-cohorts showed a homogenous distribution of T2D cases (**Supplementary Fig. 1D**). Further, rhythmicity in *alpha*-diversity, phyla proportions and taxa were confirmed in the paired sample sets (**Supplementary Fig. 1 C, E**). Similar to the cross-sectional analysis (**Fig. 3 C**), the generalized linear model was able to predict individuals at risk of developing T2D with an AUC of 0.69 (s-arOTUs) and 0.78 (s-arOTUs+BMI) (**Fig. 3 G, Supplementary Fig. 1 F**). Although T2D related differences in the relative abundance of the assigned s-arOTUs was similar between KORA and another independent twin cohort from UK (TwinsUK), the KORA-based risk signature (s-arOTUs+BMI) performed substantially worse in classifying T2D (AUC = 0.68 for pT2D) and predicting incident T2D (AUC = 0.69 for iT2D) in the regionally different TwinsUK population (**Supplementary Fig 5 A, B, Extended Data Table 3**), supporting recently published data from a Chinese cohort (12).

### Functional analysis of rhythmic microbiota and its association with T2D

We next investigated the microbiome functional changes across disease states in the predictive subcohort of KORA. The faecal metagenome of paired samples (from 2013 and 2018) from 50 KORA subjects (N = 100) was determined by shotgun sequencing. Subjects were selected based on an equal distribution of iT2D, nonT2D controls and pT2D cases (**Supplementary Table 4**). Functional pathways were identified in the shotgun sequence data using the HMP tool HUMAnN2 and annotated using KEGG. To identify those associated with diabetes, we applied a RF-based approach revealing an optimal set of 30 microbiome-encoded pathways (**Supplementary Fig. 5 C**). Models created by including only these 30 top pathways distinguished between T2D and controls with a mean AUC of 0.81, while excluding them resulted in a significant reduction of AUC to 0.60 (P < 8·10^−10^, Mann-Whitney Tests) (**Supplementary Fig. 5 C; Supplementary Table 5**). This is indicative of a strong association between these pathways and T2D, including the metabolism of amino acids (phenylalanine, cysteine, methionine, alanine, glutamine and aspartate), aromatic compounds (toluene, fluorobenzoate, chlorocyclohexane), and fatty acids (*alpha*-linoleic acid, riboflavin, drug metabolism) (**Supplementary Fig. 5 D + E**).

Twenty-six of the 30 pathways outlined above were associated (Spearman Correlation; FDR ≤ 0.1) with at least 2 of the 13 previously identified predictive s-arOTUs (**Fig. 3 H, Supplementary Table 5**). The pathways could be partitioned into two different groups based on their association patterns with the s-arOTUs. While one group of pathways (G1) was associated with bacteria related to *E. coli* (i.e. those that had significant positive associations with the clinical markers of T2D like Glucose, HOMA and Hb1AC), the other (G2) showed associations with relatives of *F. prausnitzii* (negatively correlated with clinical markers of T2D) (**Fig. 3 H**). Strong correlations were observed between *E. coli* and xenobiotic metabolism, which are also negatively associated with short-chain fatty acid biosynthesis as well as metabolism of co-factors and vitamins. The *E. coli* group of taxa was negatively associated with the *F. prausnitzii* group, which mostly occurred in combination with *C. barletti* (**Fig. 3 I)**. In contrast, *B. longum* was not associated with the presence of other taxa, but negatively correlated with xenobiotic biodegradation pathways. We next determined whether the functional associations identified could be validated in datasets from previous studies. We compared differentially abundant genes from the study by Qin et al (4) with the relative representation of the corresponding pathways in the T2D and nonT2D individuals of the KORA cohort (identified as either enriched in T2D or nonT2D). Notably, 19 of the 26 pathways could be validated in the Qin et al cohort, along with their directionality (**Fig. 3 H**). The second unique characteristic of these functional markers was their association with the arrhythmic OTUs. A recent study by Beli et al (24) investigated the microbiome changes in the diurnal cycles of experimental murine T2D and control mice using a combination of metabolomics and predictive functional profiling of amplicon sequence data. Notably, seven of the 26 pathways identified here, including xenobiotic metabolism, cysteine and methionine metabolism, *alpha-*linoleic metabolism and taurine and hypertaurine metabolism, were also reported to undergo diurnal rhythmicity in the study by Beli et al (**Figure 3 H**). Thus, the metagenomic analysis linked functions of the arrhythmic risk signature (s-arOTU) to the metabolic features of T2D (**Fig. 3 I**, **Supplementary Fig. 5 F**; **Fig. 3 I**).

## Discussion

Extending a seminal study introducing the concept of rhythmicity of the gut microbiota in two individuals (16), we here demonstrate rhythmicity in a large-scale population and, using prospective sampling, provide evidence that obesity and T2D are associated with a disruption of gut microbiota rhythmicity. We have determined a risk pattern of arrhythmic taxa which significantly contributes to the classification and prediction of T2D. A hypothesis-free machine learning strategy confirmed the predictive validity of the selected arrhythmic OTUs to discriminate between non-diabetic and T2D individuals. Together with BMI as an additional variable in the model, T2D classification significantly improved, suggesting that bacteria which become arrhythmic in T2D are indeed contributing to the risk profiling of metabolic health. Interestingly, we also detected disrupted rhythmicity in the microbiota of obese individuals. However, there was little overlap between arrhythmic OTUs of obese and diabetic individuals, indicating that BMI might be a microbiota-independent risk factor for T2D.

Development of obesity and T2D has been associated with circadian clock disruption (e.g. shift work) and gut microbiota dysbiosis (4, 7, 9, 25, 26). Despite emerging evidence for an interrelated dependency of microbiota and the host circadian system, we cannot exclude eating behaviour or the microbial communities themselves as being the key driver(s) of oscillating bacterial abundances in the gut. Nevertheless, the circadian clock seems to be required to maintain rhythmicity of microbiota composition, because oscillations are absent in mice with a genetically dysfunctional circadian clock (16). In addition, circadian signals from the microbiome affect diurnal rhythmicity of histone acetylation in intestinal epithelial cells to control metabolic responses in the host (27), supporting the hypothesis that circadian mechanisms of microbiota-host interactions contribute to metabolic homeostasis. Accordingly, loss of rhythmicity of the taxa identified in this study in T2D subjects likely results in arrhythmicity of their metabolic products. Shotgun metagenomic analysis identified 26 microbial pathways associated with xenobiotic, branch-chain amino acid, fatty acid as well as taurine metabolism, supporting a functional link between the diurnal oscillation of bacteria in the gut and metabolic homeostasis. Branched-chain amino acids were previously documented to follow circadian rhythmicity in blood samples from humans kept under 40 hours constant routine and their rhythmicity is lost in subjects with T2D (28). Although arrhythmic OTUs have been suggested to be suitable to classify and predict T2D in our regionally confined cohort, the applicability of this model to another cohort (TwinsUK) was limited, with a poor performance in sensitivity and specificity. However, 19 of the 26 pathways (73%) identified by metagenomic analysis could be validated in a regionally independent cohort of T2D individuals (3), suggesting that the functionalities of arrhythmic microbiota are relevant biomarkers. Notable among these were the pathways linked to the metabolism of certain amino acids and degradation of aromatic compounds. Among the amino acids, metabolism of alanine, aspartate, glutamate, and cysteine were positively associated with health (i.e. positive association with the health-associated taxonomic markers). The major by-products of the microbial fermentation of alanine, aspartate, glutamate and cysteine are short-chain fatty acid (SCFAs) and hydrogen sulfide (H2S), respectively (29). While the SCFAs are known for their health benefits including amelioration of insulin resistance, H2S has been especially suggested as a positive regulator of insulin sensitivity (30). In contrast, the negative association with health observed for phenylalanine metabolism was attributed to phenylethylamine and carbon dioxide (29, 30). In a similar manner p-cresol, is derived from the degradation of aromatic compounds like toluene suggested to be a negative regulator of insulin sensitivity.

Taken together, loss of diurnal oscillation in gut microbiota composition and associated rhythmic functions may contribute to the development of metabolic disorders. Whether disease-associated arrhythmic taxa and their functionality are causally linked to the metabolic phenotype of T2D remains to be studied, but these findings clearly highlight the need to monitor diurnal changes in the gut microbiome for diagnostic and prognostic investigations.

## Methods

### Longitudinal large-scale population-based cohort

All faecal samples were collected from participants of the longitudinal population-based cohort S4 KORA study (Cooperative Health Research in the Augsburg Region) in southern Germany, which started in 1999 and focuses on cardio-metabolic health, especially diabetes. The samples analyzed here are based follow-up data of the cohort in KORA-FF4 in 2013/2014 (for simplicity referred to year 2013 in the text) and KORA-Fit in 2018, respectively. Detailed study design and methods have been published previously (31). For the data analysis, 100 stool samples were excluded due to medication issues and gut-related diseases. Type-2-Diabetes and Prediabetes were defined by oral glucose tolerance test or physicians-confirmation and classified by WHO in 2013. Cases with Type-1-Diabetes were excluded.

The investigations were carried out in accordance with the Declaration of Helsinki, including written informed consent of all participants. All study methods were approved by the ethics committee of the Bavarian Chamber of Physicians, Munich (KORA-FF4 2013/14 EC No. 06068 and KORA-Fit 2018/19 EC No. 17040). The informed consent given by KORA study participants does not allow to deposit KORA data in public databases. However, data are available upon request from KORA by means of a project agreement (https://epi.helmholtz-muenchen.de/).

### Sample Collection

In 2013, faecal samples were collected from 2,076 individuals. Prospectively a subset of 699 individuals were sampled again in 2018. For the collection, participants were asked to use the received sampling kit which includes collection tubes, which are filled with 5 ml Stool stabilizer (Stratec DNA Stool Stabilizer, No. 1038111100), and to store the samples in their household refrigerator as short as possible. Participants were asked to bring the samples personally to the study centre or send it via mail. All samples were finally stored at −80°C at the study centre in Augsburg. A comprehensive data set on social-demographical characteristics, risk factors profiles, diet and medical history was ascertained amongst others.

### High-throughput 16S rRNA gene amplicon sequencing

Metagenomic DNA was isolated by a modified version of the protocol by Godon et al. (32) from 600 µl aliquots of stool mixed in DNA stabilization solution (Stratec). Briefly, microbial cells were lysed using a bead-beater with 0.1-mm glass beads (FastPrep-24 fitted with a cooling adapter). DNA was then purified on NucleoSpin gDNA columns (Machery-Nagel, No. 740230.250). DNA was either used immediately for amplicon analysis or kept frozen as aliquots of 35µl for metagenomic analysis. After DNA extraction, all pipetting steps until sequencing were conducted using a robotized liquid handler to maximize reproducibility.

PCR were conducted in duplicates. DNA was diluted in PCR-grade water (12 ng) and used as template for amplifying (25 cycles) the V3-V4 regions of 16S rRNA genes using primers 341F-ovh and 785r-ovh (33) in a two-step process shown to minimize bias (34). PCR products were pooled during cleaning using magnetic beads (Beckman Coulter). PCR-fragment concentration was determined by fluorometry and adjusted to 2nM prior to pooling. The multiplexed samples were sequenced on an Illumina HiSeq in paired-end mode (2×250 bp) using the Rapid v2 chemistry. Samples with low read counts (< 4,700) were re-sequenced on an Illumina MiSeq using v3. Control sequencings have shown no difference between both machine types (data not shown). To control for artifacts and reproducibility between runs, two negative controls (a PCR control without template DNA and a DNA extraction control consisting of 600 µl Stool stabilizer without stool) as well as a positive control using a mock community (ZymoBIOMICS, No. D6300) were included throughout for every batch of 45 samples (processed on one single 96-well-plate).

### Amplicon sequence analysis

Sequencing data was preprocessed using the IMNGS pipeline (35). Five nucleotides on the 5’ end and 3’end are trimmed for the R1 and R2 read, respectively (trim score 5) and an expected error rate of 1. Chimera were removed using UCHIME (36) and the reads of de-multiplexed samples were merged and clustered by 97% similarity using UPARSE v8.1.1861_i86 (37). OTUs occurring at a relative abundance < 0.25% across all samples were removed to prevent the analysis of spurious OTUs. Taxonomy was assigned using the RDP classifier version 2.11 and confirmed using the SILVA database (38). For phylogenetic analyses, maximum-likelihood trees were generated by FastTree based on MUSCLE alignments in MegaX (39). We used the EzBioCloud database (40) for precise identification of OTU sequences of interest.

### Validation cohort – TwinsUK

The collection of faecal samples, DNA extraction, amplification of the V4 hypervariable region of the 16S rRNA gene (primers 515F and 806R), purification and pooling were performed as previously described (41). The pooled amplicons were sequenced using the Illumina MiSeq platform with 2×250bp paired-end sequencing. The raw sequencing data was accessed via the European Nucleotide Archive (PRJEB13747). The analysis was performed on N = 1,399 individuals, including N = 46 incident Type-2-Diabetes cases as well as N = 94 Type-2-Diabetes cases that were classified using a combination of self-reported questionnaires as well as longitudinal glucose measurements.

### Statistical analysis

Statistical analysis was performed in R version 3.5.0. Absolute read counts were normalized by minimum sum counts for the calculation of within samples diversity. The contribution of covariatetes towards differences in the microbial profile of the whole cohort was determined by using multivariate permutational analysis using the R function *adonis* from the *vegan package v.2.5-6*. The explained variation of a variable is shown in R^2^ values and are considered as significant with a P-value ≤ 0.05. For the cumulative explained variation, all significant covariatetes are included in a multivariate model. Data was adjusted according to confounding factors (gender, age, BMI, physical activity, PPI, metformin, and vitamin D intake) as well as stratified according to phenotypical characteristics. Description of taxonomic composition is based on relative abundances. Between-sample diversity is calculated by generalized UniFrac using *GUniFrac* v1.1. distances. De-novo clustering is based on Ward hierarchical clustering, the selected number of clusters is chosen according to the Calinski and Harabasz index, performed with the R package *NbClust* v.3.0. For the analysis of prevalence of categorical variables between groups, a non-parametric Fisher test is used. Taxonomic differences between groups is determined by generalized linear model based on relative abundance adjusted for confounding. P-values were corrected for multiple testing using the Benjamini-Hochberg false discovery rate control procedure. Parts of this procedure have been assembled in the software pipeline Rhea (42) used for this analysis.

### Prospective Data

A subset of individuals from the cross-sectional collection from 2013 was recruited again in 2018. Stool samples from 699 individuals were collected and paired-end sequenced on an Illumina MiSeq as described above. T2D was classified based on HbA1c value (%) an incident T2D case is defined as HbA1c < 6.5% in 2013 and HbA1c ≥ 6.5% in 2018 because oral glucose tolerance test was not available in 2018. Sequencing data from the sub-cohort of paired subjects is used to predict T2D. Subjects with unclassified T2D status are excluded. To avoid bias due to overfitting, the 699 samples from 2013, which are part of the 2018 follow-up dataset, are retained for independent validation.

### Illustration of diurnal profiles

A high-resolution time course was generated by merging samples taken between 5:00 am and 24:00 pm in two-hour intervals, and a larger 4-hour interval at night from 00:01 am to 4:59 am to compensate for the low sample size in some groups during the night time points. The merged data points are illustrated using GraphPad Prism v6.01 (GraphPad Software) with the sample size of every groups per time point indicated below the data point in the individual graphs. To demonstrate the overall phase relationship and periodicity of all OTUs together, heatmaps have been generated using the online tool *Heatmapper* (http://www.heatmapper.ca; (43)). The raw data of each OTU were merged in the above indicated intervals, sorted by the peak phase based on cosine-wave regression analysis (described below) and scaled in each row according to the highest abundance of the OTU.

### Diurnal analysis of microbiome data sets

Statistical analyses were conducted with GraphPad Prism v6.01 (GraphPad Software) and the R script JTK_CYCLE v3.1.R (44) using Rstudio v1.1.456 (Rstudio Inc.). To efficiently identify and characterize diurnal oscillations in large datasets, circadian variation was tested by fitting a cosine-wave equation: y = baseline+(amplitude·cos(2·π·((x−[phase shift)/24))) or a double harmonic cosine-wave equation: y = baseline+([amplitude A]·cos(2·π·((x−[phase shift A])/24))) + ([amplitude B]·cos(4·π·((x−[phase shift B])/24))) on *alpha*-diversity and relative abundance, with a fixed 24-h period. The goodness of fit was corrected for multiple comparisons and the significance was determined using an F-test. Results from the cosine- and harmonic cosine-wave regression were compared with a widely used rhythmicity detection algorithms JTK_CYCLE (**Supplementary Fig. 1**), which employs a non-parametric algorithm detecting sinusoidal signals (44), whereby JTK presents the highest false negative rates (45). Each P-value was Bonferroni-adjusted for multiple testing. A statistically significant difference was assumed when P ≤ 0.05. The high-density time sampling allowed the identification of diurnal-regulated microbiota with a high statistical power (45).

### Detection of (ar)rhythmic OTUs (arOTUs) and random-forest selected arrhythmic OTUs (s-arOTUs)

The relative abundance of each OTU was assessed for a 24-h rhythmicity using the cosine-wave regression. On total, 87 OTUs showed diurnal fluctuation in subjects with nonT2D, Prediabetes or a BMI < 30, whereas no significant rhythmicity was detected in subjects with T2D or BMI ≥ 30 for those OTUs (**Fig. 2H, Supplementary Table 2**). These 87 OTUs were further analyzed for differential 24-h time-of-day patterns using the Detection of Differential Rhythmicity (DODR) R packages (46). Resulting DODR P-values were corrected for multiple comparisons and at the corrected P ≤ 0.05 significance level DODR detected 26 OTUs, referring to as arOTUs, with significantly different 24-h time-of-day patterns when comparing nonT2D with T2D, prediabetes with T2D and BMI < 30 with BM ≥ 30 (Fig. 3F). An overlap of these 26 arOTUs with the previously defined 30 OTUs (abundant OTUs), which showed a significantly different averaged abundance between nonT2D and T2D (**Fig. 1J, Extended Data Table 2**), identified 13 OTUs. The abundance of these 13 OTUs followed a 24-h rhythm in the control groups (nonT2D, BMI < 30), but were arrhythmic in T2D and BMI ≥ 30 and in addition significantly changed their abundance (s-arOTUs).

### Classification model

A machine-learning algorithm is applied to the cross-sectional data (n = 1,340) from 2013 to classify T2D, excluding the paired-subcohort (year 2013 = 699). Further, the most prevalent (> 10%) and abundant (> 0.1%) OTUs are selected for the analysis, resulting in 425 OTUs. A random forest model was used to classify binary outcome variables based on a combination of BMI and microbial composition with a 5-fold cross validation by using *randomForest* from the R package *randomForest* v4.6-14. To receive a robust and generalizable classification model, the machine-learning algorithm was applied 100-times iteratively assigning randomly individuals to either the training (80%) or test test set (20%). For the training set, a subset of equally distributed T2D and nonT2D cases was taken to train the model. The model was then validated on the 20% test set. Based on out-of-bag error rates and Gini index, the most important features were selected for each iteration using *rfcv* from R package *randomForest* v4.6-14. Features which appeared in at least 50% of all 100 random forest models were considered as classification feature for the final model.

### Prognostic model for T2D

For the risk prediction of T2D, a generalized linear model for binomial distribution and binary outcome (logit) was generated using the previously selected features based on arrhythmic OTUs including BMI as additional variable. For the model, a generalized linear model was generated on the cross-sectional cohort excluding the paired sub-cohort. To verify the importance of the selected features, a generalized linear model for control OTUs (rndOTUs, equal number of OTUs as in s-arOTUs) are implemented repetitively 100-times. The randomly selected control OTUs (rndOTUs) neither show rhythmicity in disease nor non-disease stages.

To apply the generated models to the unknown data from the paired sub-cohort of both time points, a blast search was performed assigning sequences of the selected features to the corresponding sequences of the new dataset. Sequences with an identify of 97%, coverage of 80% and an E value below 10^−5^ are considered as a hit. If there is no matching sequence available when using the above thresholds, the best match is taken instead. Sequences are uniquely assigned to a reference sequence. For the prediction of T2D, the relative OTU abundance of year 2013 is considered as baseline. Individuals at this stage are not classified as T2D. The two endpoints are incident T2D cases and nonT2D.

### Metagenomic data selection

From of the prospective data, we chose a subset of 100 paired individuals showing interesting phenotypic characteristics for shotgun sequencing and metabolomic analysis. The dataset includes incident T2D cases, T2D cases and nonT2D controls, which are otherwise metabolically healthy. Shotgun sequencing of the isolated DNA from stool samples, as well as the analysis, were conducted by the APC Microbiome Institute, Cork (Ireland).

The taxonomic and functional annotation of the shotgun feacal metagenomic datasets were performed using the *metaphlan2* (47) and HUMAnN2 (48) pipelines. Gene families detected using the HUMAnN2 approach were then mapped to the KEGG pathway orthology (49) scheme using internal mappings within HUMAnN2. *Psych* R package v1.8.12 was used to compute the correlation between the OTU markers with the clinical markers and the KEGG pathways (Spearman correlation filtered with Benjamini-Hochberg corrected FDR ≤ 0.1). For identifying the top disease-predictive pathways, we used an iterative random forest approach *randomForest* from R package *randomForest* v4.6-14, where we performed 100 iterations, each time taking 50% of the samples with T2D (from both 2013 and 2018) and an equal number of controls and tested the same model on the remaining 50%, again with an equal number of controls. Mean AUC and mean feature importance scores were then computed across iterations using standard R functions. For the validation in the cohort of Qin et al. (4) cohort, the pathway affiliations of the differentially abundant KEGG orthologues identified in this study were obtained. The relative representation of the marker pathways in the T2D-enriched and control-enriched were then computed. Pathways that were only detected in a group or had more than two-fold increase of representation in the given group were identified as enriched in that group (T2D or nonT2D), respectively.

## ADDITIONAL INFORMATION (CONTAINING SUPPLEMENTARY INFORMATION LINE (IF ANY) AND CORRESPONDING AUTHOR LINE)

### ACKNOWLEDGEMENT

The KORA study was initiated and financed by the Helmholtz Zentrum München – German Research Center for Environmental Health, which is funded by the German Federal Ministry of Education and Research (BMBF) and by the State of Bavaria. TwinsUK is funded by the Wellcome Trust, Medical Research Council, European Union, Chronic Disease Research Foundation (CDRF) and the National Institute for Health Research (NIHR)-funded BioResource, Clinical Research Facility and Biomedical Research Centre based at Guy’s and St Thomas’ NHS Foundation Trust in partnership with King’s College London. The Technical University of Munich provided funding for the ZIEL Institute for Food & Health. Caroline Ziegler and Angela Sachsenhauser provided outstanding technical support for sample preparation and 16S rRNA gene amplicon sequencing.

### AUTHOR CONTRIBUTIONS

DH conceived and coordinated the project. SR, SK, TC performed 16S rRNA gene sequencing data analysis. ELA and TSG performed shotgun metagenomics and data analysis. SR and ML performed informatics analysis. DH, TC, POT, JB supervised the work and data analysis. KN supported sample preparation and 16S rRNA gene sequencing analysis. AF provided help for sample analysis. HG, MT, WR and AP provided KORA data and performed sample collection. CILR, JTB and TS provided TwinsUK data. SR, SK, TC, ML, TSG, POT and DH wrote the manuscript. All authors reviewed the manuscript.

Sandra Reitmeier, Silke Kießling and Thomas Clavel contributed equally to this work.

### CORRESPONDING AUTHOR

Correspondence to Prof Dr. Dirk Haller, Chair of Nutrition and Immunology, Technical University of Munich, Gregor-Mendel-Str. 2, 85354 Freising, Germany

**Supplementary Figure 1.**
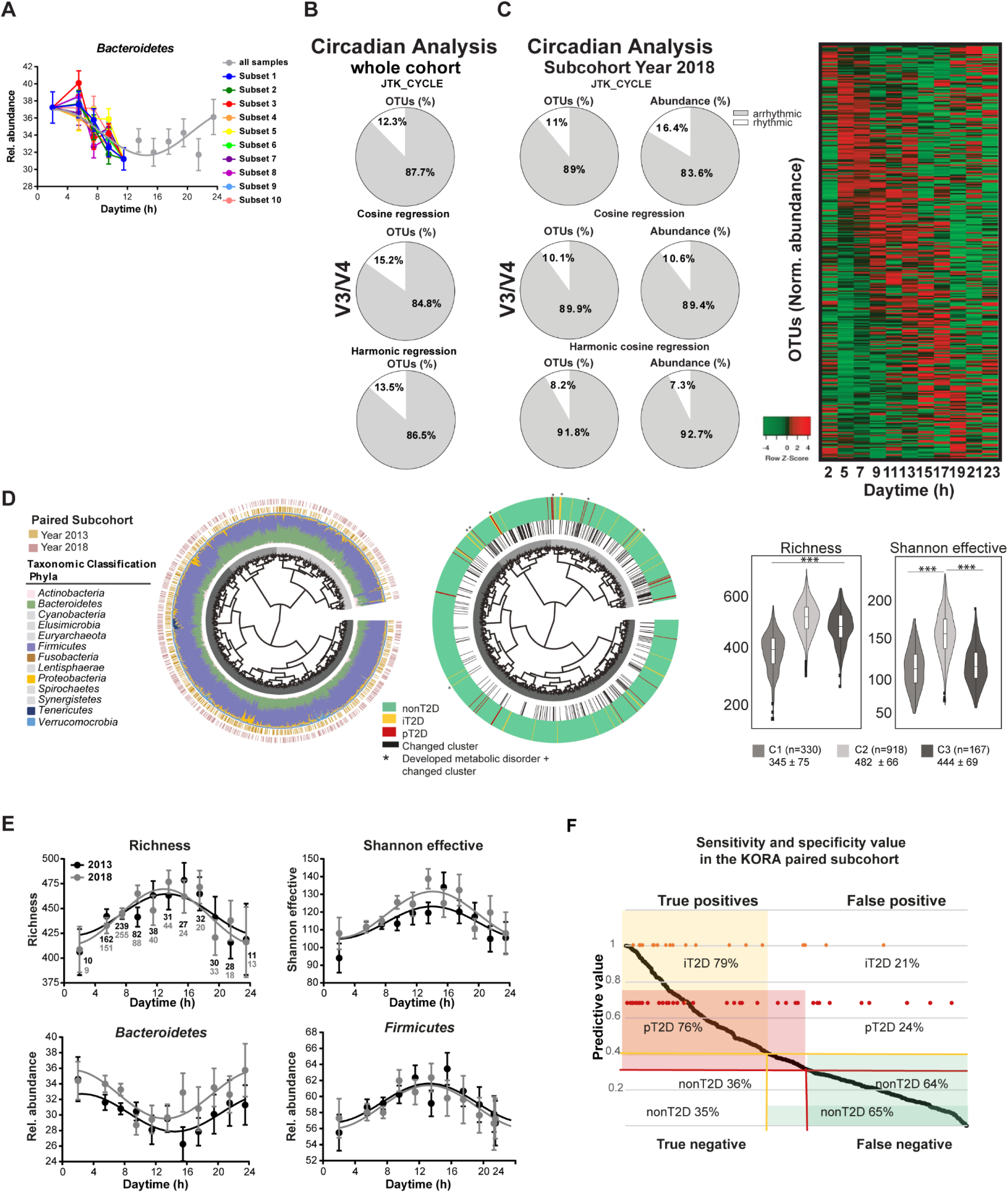
Diurnal rhythm and microbiota profiling of KORA in year 2013 and 2018. **A:** Circadian analysis of rOTUs in % based on JTK_CYCLE, cosine-wave regression and harmonic cosine-wave regression of the subcohort (year 2018). Heatmap depicting the overall phase relationship and periodicity of OTUs is generated by the raw data of all 425 OTUs (relative abundance: 0.1%, prevalence: 10%) ordered by phase and normalized to the peak of each OTU. **B:** Circadian analysis of rOTUs in % based on JTK_CYCLE, cosine-wave regression and harmonic cosine-wave regression of the whole cohort. **C:** Circadian profiling of the relative abundance of *Bacteroidetes* based on 10 different randomly selected subsets with equal sample size (N = 25) between 5 and 11am. **D:** Phylogenetic tree of 1,401 paired individuals from year 2013 and year 2018 represented by leaves. Subjects are grouped according to similar microbial profile calculated by generalized UniFrac distance. Based on unsupervised hierarchical clustering, individuals are assigned into three cluster C1 = 330, C2 = 918 and C3 = 167 (greyish colored leaves). Taxonomic composition on phyla level for each subject is shown as colored stacked barplots around the circle. Color strips around the circle are referring to the sample collection year. Marked leaves of the right phylogenetic tree indicate a cluster change and the T2D status. *Alpha*-diversity is represented by the three cluster with significant differences in richness and Shannon Effective (P ≤ 1.0^e-5^, P ≤ 1.0^e-5^). **E:** Diurnal profile of the relative abundance of *alpha*-diversity and the phyla of subjects from the KORA subcohort 2013 (N = 699, black) or subcohort 2018 (N = 699, grey). Significant rhythms are illustrated with fitted cosine-wave curves (cosine-wave regression, P ≤ 0.05). **F:** Shows the predictive values for the matched KORA cohort. Yellow circles in the upper part show incident T2D cases (iT2D) and red circles in the lower part show the classification of persisting T2D cases (pT2D). Different threshold for prediction (0.3) and classification (0.4) of T2D are shown in the graph, with the aim to find the best balance between sensitivity and specificity.

**Supplementary Figure 2.**
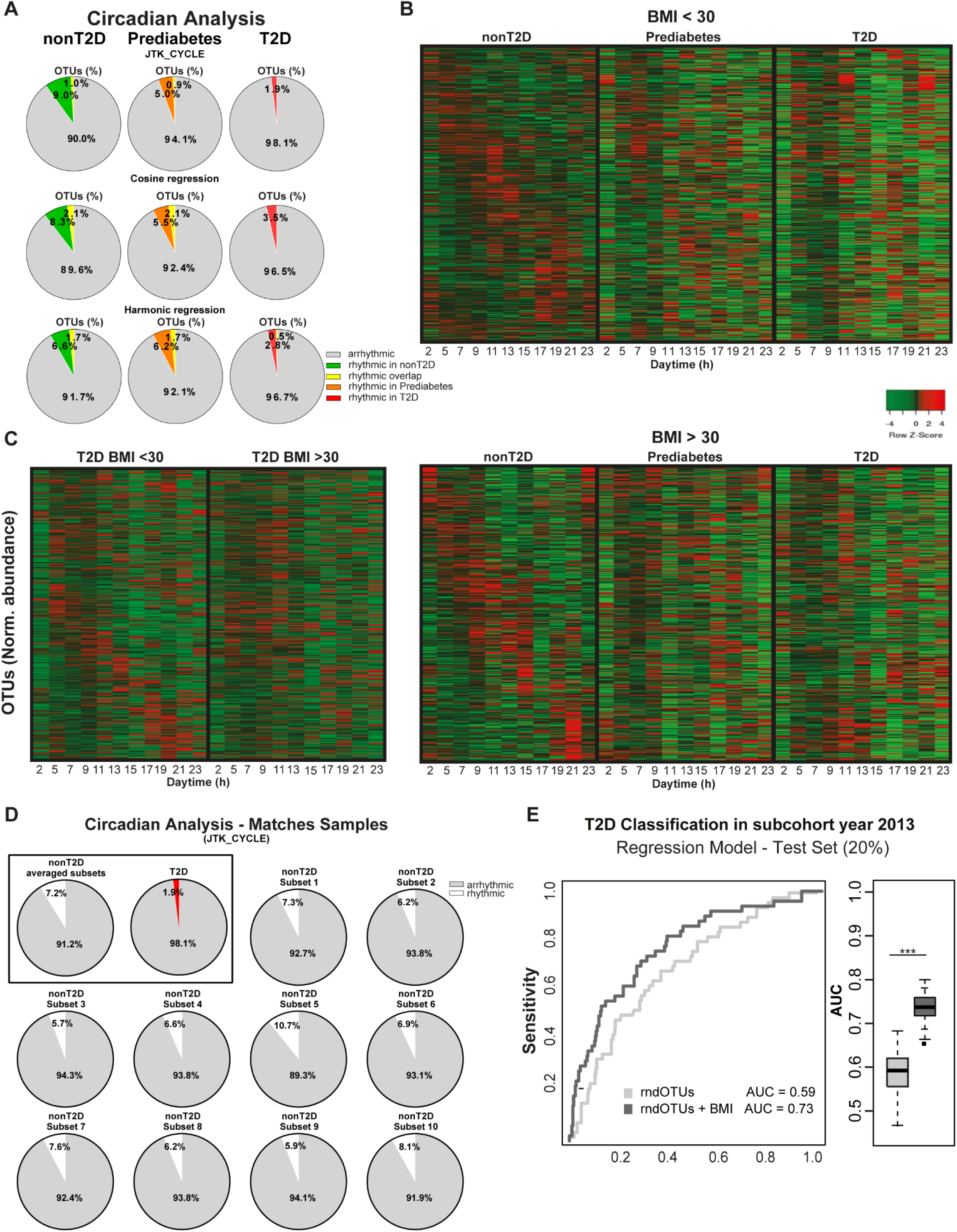
Microbial rhythmicity in Type-2-Diabetes. **A:** Circadian analysis of rhythmic (colored) and arrhythmic (grey) clustered OTUs and their relative abundance in percent based on JTK_CYCLE, cosine-wave regression and harmonic cosine-wave regression in nonT2D (left, N = 1,255, green), prediabetes (middle, N = 352, orange) and T2D (right, N = 269, red). **B:** Heatmap depicting the overall phase relationship and periodicity of OTUs is generated by the raw data of all 422 OTUs found with sufficient prevalence and abundance ordered by phase according to nonT2D subjects with either a BMI < 30 (top) or with a BMI ≥ 30 (bottom) and normalized to the peak of each OTU. **C:** Heatmap showing the normalized daytime dependent abundance of OTUs generated by the raw data of all 422 OTUs (relative abundance: 0.1%, prevalence: 10%) and ordered by the peak phase according to subjects with T2D and with a BMI < 30 normalized to the peak of each OTU. **D:** Circadian analysis of rOTUs in % based on JTK_CYCLE, cosine-wave regression and harmonic cosine-wave regression of the samples from nonT2D matched to the sample size of T2D for every sampling time point between 5 and 11am. **E:** ROC curve for classification of T2D in an independent test set. Generalized linear model is based on 13 randomly selected control OTUs +/− BMI (rndOTUs, grey curves). The iterative calculated AUCs are shown by boxplots and are significantly different between the models.

**Supplementary Figure 3.**
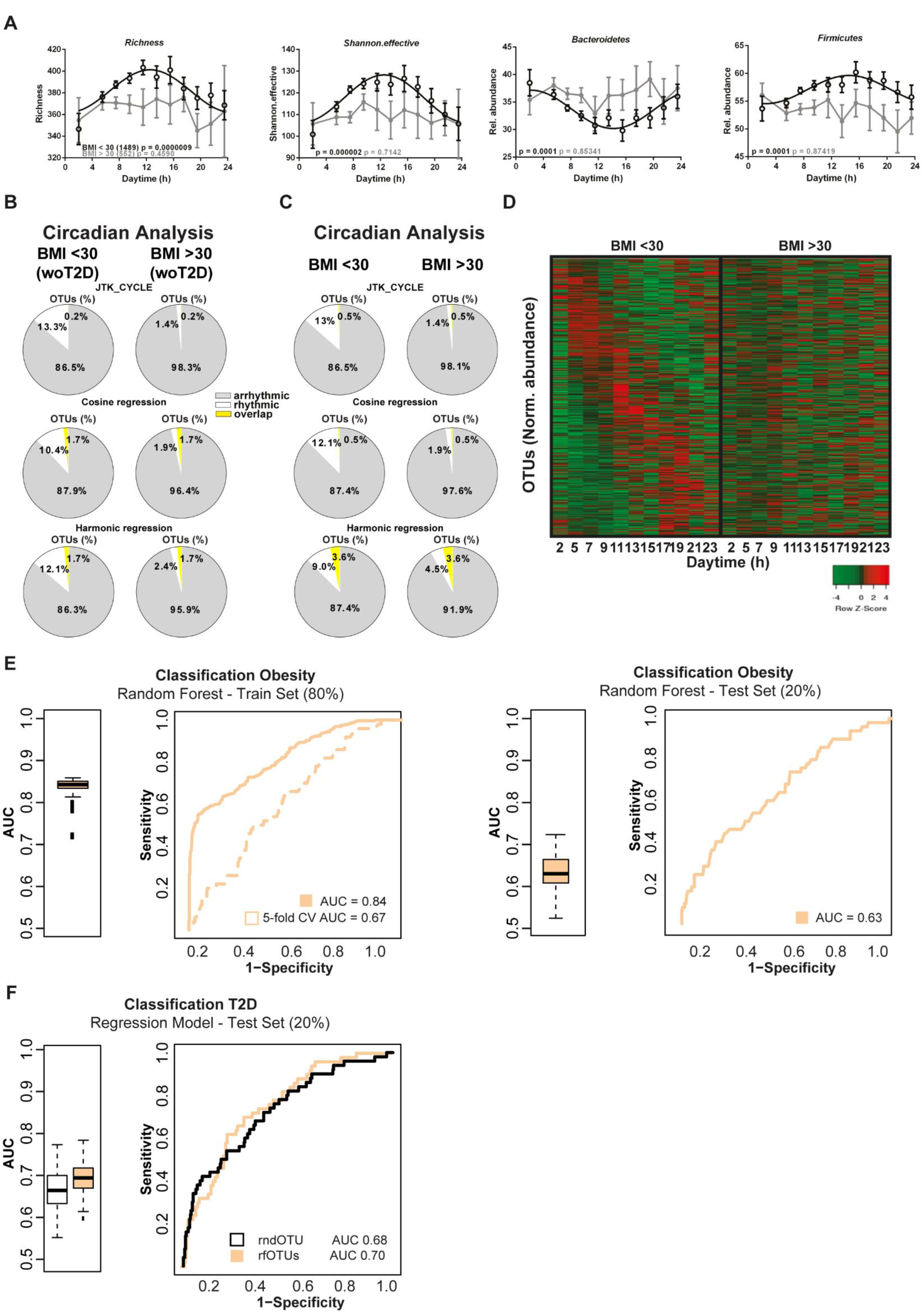
Diurnal rhythmicity in obesity and its association with Type-2-Diabetes. **A:** Diurnal profile of the relative abundance of *alpha*-diversity and the phyla of subjects (with either a BMI < 30 (N = 1,489, black) or with a BMI ≥ 30 (N = 552, grey). Significant rhythms are illustrated with fitted cosine-wave curves, otherwise data are simply connected by straight lines between data points, indicating no significant cosine fit curves (cosine-wave regression, P ≤ 0.05). **B-C:** Circadian analysis of rOTUs in % based on JTK_CYCLE, cosine-wave regression and harmonic cosine-wave regression of individuals with (**B**) a BMI < 30 or ≥ 30, but without (wo.) T2D and (**C**) a BMI < 30 or ≥ 30. **D:** Heatmap showing the normalized daytime dependent abundance of OTUs generated by the raw data of all 422 OTUs (relative abundance: 0.1, prevalence: 10%) and ordered by the peak phase according to subjects with diabetes and a BMI < 30 normalized to the peak of each OTU. **E:** ROC curve of random forest trained model for classification of individuals with BMI ≥ 30 and BMI < 30 with mean AUC for the training set and 5-fold cross validation. Iterative AUC values are shown in the left boxplot. Right ROC curve represents the results out of random forest model for an independent validation set. **F:** ROC curve and AUC of generalized linear model for selected features from **E** to classify T2D based on a BMI – related OTU list (orange) as well as randomly selected OTUs as control list (black).

**Supplementary Figure 4.**
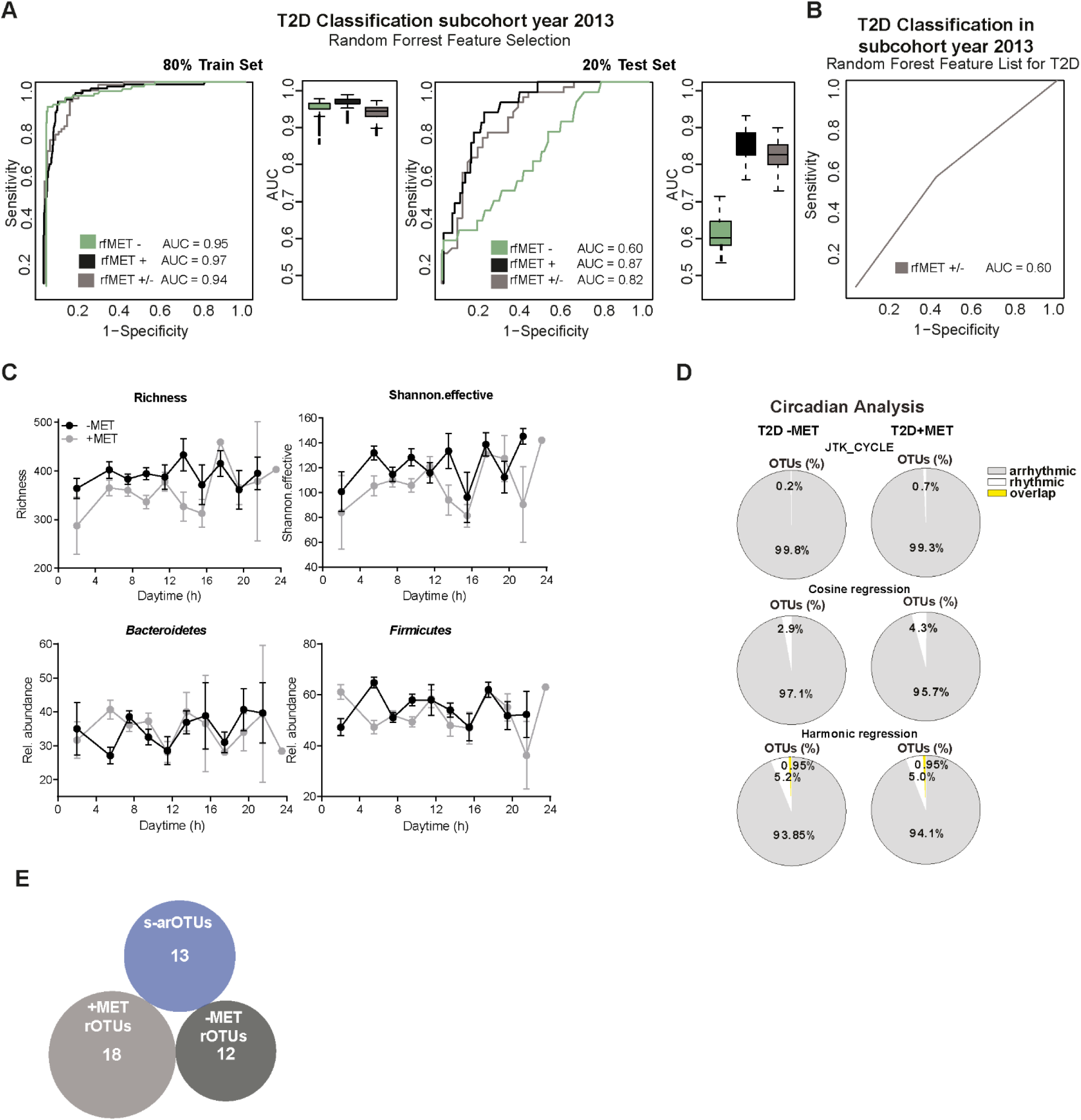
Influence of metformin in classification of Type-2-Diabetes and its impact towards diurnal rhythmicity. **A:** ROC curve of random forest to evaluate the effect of metformin intake in T2D. Individuals are stratified in three groups, T2D+MET compared to nonT2D compared to nonT2D (black), T2D-MET (green) and T2D +/− metformin (grey). In case of the last group, classification is based on the outcome metformin. Iterative AUCs are represented in boxplots for training and independent test sets as well as the mean AUC of all groups for training and independent test. **B:** ROC curve of the results based on generalized linear regression model for thirteen selected arrhythmic OTUs (compare Fig. 3 **D**, s-arOTU + BMI) for differentiation of metformin in T2D. **C:** Diurnal profile of the relative abundance of *alpha*-diversity indicated by richness and Shannon effective and the phyla, *Bacteroidetes* and *Firmicutes* of T2D subjects which take metformin (+MET, grey, N = 132) or not (-MET, black, N = 136). Data point were connected by straight lines to illustrate no significant rhythm based on a fitted cosine-wave curve (cosine-wave regression, P > 0.05). **D:** Circadian analysis of rhythmic (white) and arrhythmic (grey) OTUs and their relative abundance in percent based on JTK_CYCLE, cosine-wave regression and harmonic cosine-wave regression in T2D-MET (left) and T2D+MET (right). **E:** Venn diagram to illustrate no overlap between the s-arOTUs and all identified rOTUs in either T2D–MET or T2D+MET based on cosine-wave regression analysis (P ≤ 0.05).

**Supplementary Figure 5.**
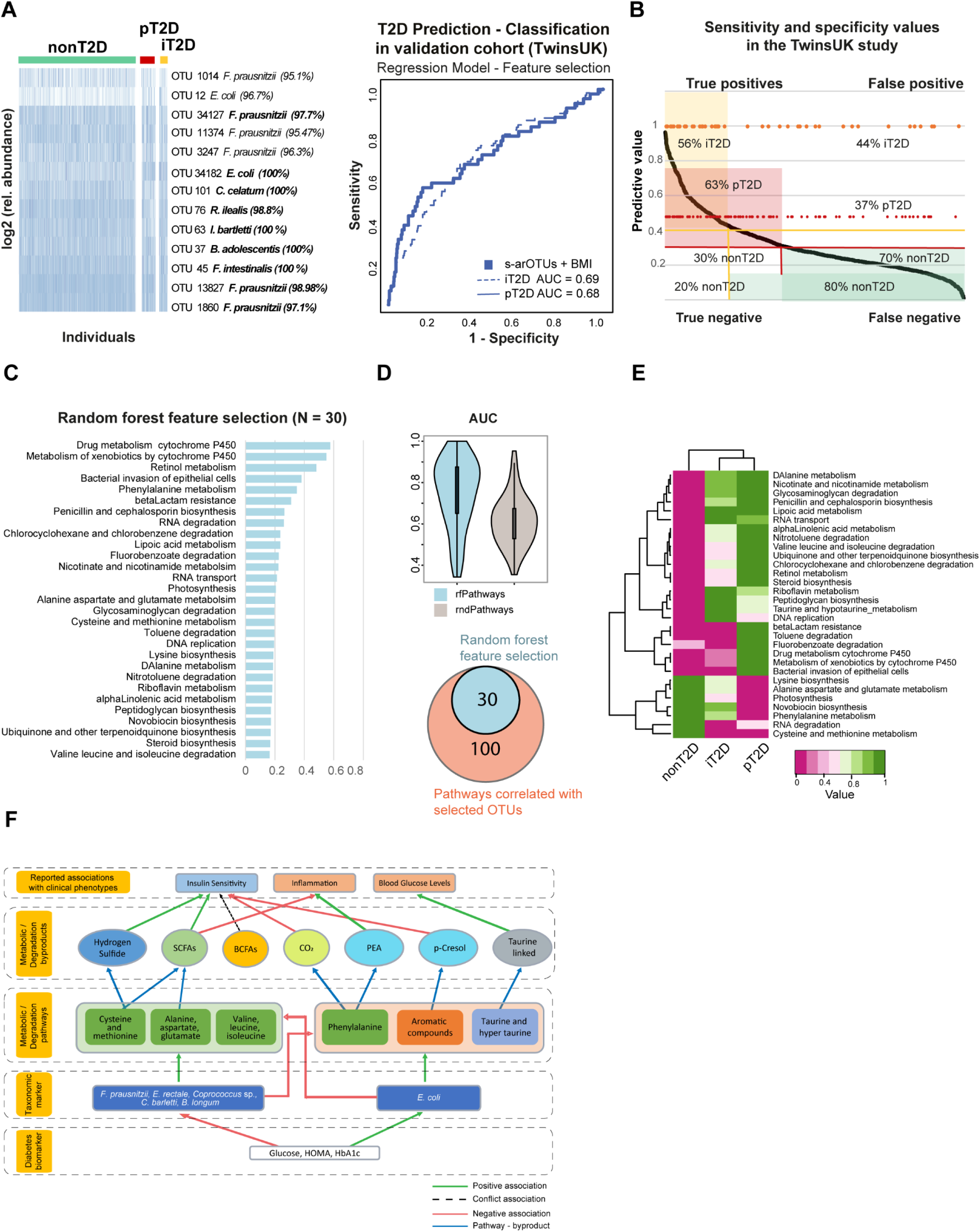
Validation and functional association of arrhythmic risk signature in association with Type-2-Diabetes. **A: Left panel**, Heatmap showing the log-transformed relative abundance of the TwinsUK OTUs (y-axis) identified as representatives of the a-srOTUs (Supplemenatry Table 3). Individuals (x-axis) are grouped according to their T2D status. Taxonomic classification of OTUs is shown by their species names and 16S rRNA sequence similarity (%), bold indicates ≥ 97% similarity. **Right panel**, Classification (pT2D, straight blue line, AUC = 0.69) and predictive analysis of T2D (iT2D, dashed blue line, AUC = 0.68) in independent TwinsUK cohort. The prediction of incident T2D cases (iT2D) is based on the 13 s-arOTUs +/− BMI. **B**: Shows the predictive values of the validation cohort TwinsUK. Yellow circles in the upper part show for incident T2D cases (iT2D) and red circles in the lower part show the classification of persisting T2D cases (pT2D). Different threshold for prediction (0.3) and classification (0.4) of T2D are shown in the graph, with the aim to find the best balance between sensitivity and specificity. **C: Left panel**, List of top 30 optimal marker pathways along with their feature importance score. The feature importance scores increased sharply for the top 30 pathways and decreased linearly for all pathways after the top 30. Consequently, the top 30 pathways were identified as the optimal set of functional markers. **Right panel**, Comparison of AUC distribution of iterative random forest models obtained by including only the top 30 pathways and excluding the top 30 pathways. Models incorporating only the top 30 pathways could predict T2D individuals from controls with a mean AUC of 0.81. Excluding these pathways resulted in a significant decrease (P < 3·10^−9^) to mean AUCs around 0.6. **D:** In total 130 KEGG pathways are significantly associated with at least one of the selected OTUs. Comparing the random forest selected KEGG pathways with the risk signature associated pathways, shows a 100% overlap. **E:** Heatmap shows the associated of KEGG pathways and T2D, by stratifying the individuals into nonT2D, iT2D and T2D it is shown that some of the pathways are associated with nonT2D while other are more prevalent in T2D. **F:** A schematic flow Integrating T2D signature, rhythmicity and microbiome function in the metagenomes. The flow correlates the major disease-associated metabolic pathways identified in the current study, their associations with the disease/health-associated taxonomic markers, the metabolic byproducts originating from these pathways, and the previously known associations of these byproducts with the various clinical phenotypes. The green-arrows are positive association, red is negative, conflicting associations are as dashed arrow (BCFA has been shown to be positively associated insulin sensitivity in some paper, while others show no association). Blue-arrows link pathways to their metabolic by-products.

**Supplementary Table 1.** Descriptive overview diabetes associated characteristics of the KORA cohort (year 2013). Shown as mean values ± standard deviation. Significance is calculated by Kruskal-Wallis one-way ANOVA for numerical data and Fisher test for categorical data.

|  | non Type-2-Diabetes<br>(N=1270) | Prediabetes<br>(N=356) | Type-2-Diabetes<br>(N=277) | p-value |
| --- | --- | --- | --- | --- |
| Male (%) | 566 (44.56%) | 202 (56.74%) | 164 (59.21%) | 1.65E-07 |
| Age | 57 ± 11 | 65 ± 11 | 69 ± 9 | < 2.2e-16 |
| Waist (cm) | 93 ± 13.09 | 103.06 ± 12.50 | 107.34 ± 13.13 | 1.16E-09 |
| Hip (cm) | 105.21 ± 8.89 | 109.15 ± 9.21 | 110.72 ± 10.63 | < 2.2e-16 |
| BodyFat (%) | 31.89 ± 7.04 | 34.51 ± 6.59 | 35.37 ± 7.03 | 3.98E-01 |
| BMI | 26.62 ± 4.46 | 29.75 ± 4.68 | 31.09 ± 5.38 | 9.92E-03 |
| Triyglyceride (mg/dl) | 109.77 ± 66.51 | 141.21 ± 70.81 | 154.89 ± 84.77 | 1.58E-06 |
| HbA1c (%) | 5.32 ± 0.32 | 5.60 ± 0.33 | 6.49 ± 1.03 | < 2.2e-16 |
| HOMA-IR | 2.09 ± 1.24 | 3.85 ± 2.26 | 5.48 ± 3.72 | < 2.2e-16 |
| Glucose (mg/dl) | 94.25 ± 7.32 | 106.40 ± 9.72 | 136.79 ± 34.30 | < 2.2e-16 |
| Glucose (mg/dl) after OGTT | 97.20 ± 20.09 | 148.13 ± 27.30 | 222.27 ± 65.88 | < 2.2e-16 |
| Insulin (pmol/l) | 53.33 ± 29.54 | 87.43 ± 49.65 | 107.80 ± 93.70 | < 2.2e-16 |
| Insulin (pmol/l) after OGTT | 283.55 ± 245.85 | 653 ± 496.19 | 794.53 ± 491.21 | < 2.2e-16 |
| HDL-Cholsterin (mg/dl) | 69.48 ± 18.65 | 61.34 ± 18.11 | 57.84 ± 16.39 | 9.76E-06 |
| LDL-Cholesterin (mg/dl) | 135.49 ± 34.96 | 139.03 ± 36.69 | 128.32 ± 37.32 | 3.32E-01 |
| Vitamin D (ng/ml) | 25.87 ± 11.98 | 23.65 ± 11.89 | 22.62 ± 12.37 | 7.79E-01 |
| Metformin | 0 | 0 | 142 (48.73%) | < 2.2e-16 |
| PPI | 99 (7.80%) | 56 (15.73%) | 48 (17.32%) | 3.92E-07 |

**Supplementary Table 2.** Sequencing depth of the 16S rRNA gene sequencing. Tables shows the number of generated reads for both years (2013 and 2018). Samples with too low sequencing quality (< 4,700) were resequenced and included in the final dataset. Last row shows the number of generated OTUs.

|  | 16S rRNA gene sequencing<br>2013/2014 | 16S rRNA gene sequencing<br>2018 |
| --- | --- | --- |
| Samples wo resequencing | 2,186 | 900 |
| Average no. reads | 87,655 | 29,880 |
| Max. reads | 211,648 | 60,141 |
| Min reads | 16 | 7,363 |
| Sum all reads | 187,144,068 | 26,892,897 |
| Samples resequenced | 207 | not resequenced |
| Average no. reads | 145,328 |  |
| Max reads | 581,380 |  |
| Min reads | 36,164 |  |
| Sum all reads | 300,083,040 |  |
| Samples incl. resequencing** | 2,137 | 900 |
| Average no. reads | 99,386 | 29,880 |
| Max reads | 581,380 | 60,141 |
| Min reads | 36,164 | 7,363 |
| Sum all reads | 212,289,588 | 26,892,897 |
| Number OTUs | 2,089 | 1,695 |

\*\* removing samples with low quality

**Supplementary Table 3.** Overview of subjects include in the analysis. The table shows the number of subjects (N) and OTUs for each analysis. Due to different issues of each analysis, the exclusion and inclusion criteria are customized (see Methods).

|  | Year | N sequenced | N after exclusion | No. OTUs analysed |
| --- | --- | --- | --- | --- |
| Cross-sectional analysis | 2013/2014 | 2,137 | 1,976 | 422 |
| Classification model | 2013/2014 | 2,137 | 1,340 | 425 |
| Cross-sectional analysis | 2018 | 900 | 699 | 318 |
| Prediction model | 2013/2014 | 2,137 | 699* | 2,275** |
|  | 2018 | 900 | 699 |  |

\* different exclusion criteria \*\* no OTU filtering for BLAST search

**Supplementary Table 4.** Samples selected for metagenomic shotgun sequencing. Overview of the 50 selected paired samples (from year 2013 and 2018) for metagenomic shotgun sequencing. Subjects are selected due to Diabetes status. A nonT2D control was selected based on comparable distribution of age and gender as well as an equal distribution of obese individuals. All incident T2D cases are selected for metagenomic sequencing.

| | Year | no. | mean Age | Gender | BMI | Mean ± SD HbA1c |
| --- | --- | --- | --- | --- | --- | --- |
| nonT2D | 2013 | 8 | 57 | 50% male | 50% Obese | 5.41 ± 0.31 |
| | 2018 | 8 | 62 | | 50% Obese | 5.42 ± 0.30 |
| incident T2D | 2013 | 20 | 59 | 60% male | | 6.06 ± 0.33 |
| | 2018 | 20 | 64 | | | 7.10 ± 0.86 |
| remained T2D | 2013 | 14 | 62 | 47% male | | 7.64 ± 1.19 |
| | 2018 | 14 | 67 | | | 7.75 ± 1.19 |
| Prediabetes | 2013 | 8 | 59 | 50% male | 50% Obese | 5.33 ± 0.35 |
| | 2018 | 8 | 64 | | 50% Obese | 5.755 ± 0.36 |
| TOTAL |  | 100 |  |  |  |  |

**Supplementary Table 5.** Functional pathways in association with Type-2-Diabetes. The shown pathways (N = 30) are selected by random forest. Rows are ordered due to significance and coloured pathways (blue) are significantly different between T2D and nonT2D. Disease column refers to increased mean value in either T2D or nonT2D.

|  | Mann-Whitney U P-value | Disease | FDR Corrected P-value | Mean in T2D | Mean in nonT2D |
| --- | --- | --- | --- | --- | --- |
| Retinol metabolism | 0.003 | T2D | 0.094 | 0.0002 | 0.0001 |
| Drug metabolism cytochrome P450 | 0.005 | T2D | 0.156 | 0.0002 | 0.0002 |
| Metabolism of xenobiotics by cytochrome P450 | 0.005 | T2D | 0.156 | 0.0002 | 0.0002 |
| Bacterial invasion of epithelial cells | 0.014 | T2D | 0.373 | 0.0000 | 0.0000 |
| Penicillin and cephalosporin biosynthesis | 0.018 | T2D | 0.466 | 0.0001 | 0.0000 |
| betaLactam resistance | 0.020 | T2D | 0.511 | 0.0004 | 0.0003 |
| Lipoic acid metabolism | 0.020 | T2D | 0.511 | 0.0012 | 0.0009 |
| Fluorobenzoate degradation | 0.023 | T2D | 0.533 | 0.0001 | 0.0001 |
| Lysine biosynthesis | 0.025 | nonT2D | 0.550 | 0.0141 | 0.0147 |
| DAlanine metabolism | 0.046 | T2D | 0.960 | 0.0018 | 0.0016 |
| Alanine aspartate and glutamate metabolism | 0.047 | nonT2D | 0.960 | 0.0210 | 0.0219 |
| Ubiquinone and other terpenoidquinone biosynthesis | 0.049 | T2D | 0.960 | 0.0032 | 0.0026 |
| Phenylalanine metabolism | 0.049 | nonT2D | 1 | 0.0030 | 0.0034 |
| Riboflavin metabolism | 0.063 | T2D | 1 | 0.0041 | 0.0038 |
| alphaLinolenic acid metabolism | 0.063 | T2D | 1 | 0.0002 | 0.0001 |
| Photosynthesis | 0.067 | nonT2D | 1 | 0.0101 | 0.0108 |
| Valine leucine and isoleucine degradation | 0.072 | T2D | 1 | 0.0035 | 0.0032 |
| Steroid biosynthesis | 0.076 | T2D | 1 | 0.0001 | 0.0000 |
| Cysteine and methionine metabolism | 0.082 | nonT2D | 1 | 0.0175 | 0.0180 |
| Glycosaminoglycan degradation | 0.084 | T2D | 1 | 0.0014 | 0.0011 |
| Nitrotoluene degradation | 0.087 | T2D | 1 | 0.0009 | 0.0007 |
| Nicotinate and nicotinamide metabolism | 0.119 | T2D | 1 | 0.0083 | 0.0080 |
| Novobiocin biosynthesis | 0.119 | nonT2D | 1 | 0.0030 | 0.0031 |
| RNA transport | 0.122 | T2D | 1 | 0.0014 | 0.0013 |
| Toluene degradation | 0.163 | T2D | 1 | 0.0027 | 0.0024 |
| Chlorocyclohexane and chlorobenzene degradation | 0.175 | T2D | 1 | 0.0001 | 0.0001 |
| RNA degradation | 0.202 | nonT2D | 1 | 0.0068 | 0.0069 |
| Peptidoglycan biosynthesis | 0.212 | T2D | 1 | 0.0122 | 0.0119 |
| DNA replication | 0.327 | T2D | 1 | 0.0087 | 0.0087 |
| Taurine and hypotaurine metabolism | 0.551 | T2D | 1 | 0.0031 | 0.0030 |

